# Apico-basal cell compression regulates Lamin A/C levels in Epithelial tissues

**DOI:** 10.1101/2020.05.18.102509

**Authors:** K Venkatesan Iyer, Natalie A. Dye, Suzanne Eaton, Frank Jülicher

## Abstract

Nuclear lamina bridges mechanical forces from the cytoskeleton to the nucleus, to initiate nuclear mechanotransduction. The concentration of nuclear Lamin proteins, particularly Lamin A/C is crucial for the mechanical properties of the nucleus and nuclear mechanotransduction. Recent studies in mesenchymal tissues show that the concentration of Lamin A/C scales with stiffness and concentration of the underlying extracellular matrix (ECM). But in epithelial tissues, that lack a strong cell-ECM interaction, it is still unclear how Lamin A/C is regulated. Here, we show that concentration of Lamin A/C in epithelial tissues scales with apico-basal compression of cells and is independent of ECM concentration. But, ectopically altering the concentration of Lamin A/C does not influence cell shapes in epithelial tissues. Using genetic perturbations in *Drosophila* epithelial tissues, we reveal that apico-basal cell compression regulates the concentration of Lamin A/C by deforming the nucleus. We observe a similar mechanism of Lamin A/C regulation in mammalian Madin Darby Canine Kidney (MDCK) cells suggesting that this mechanism is evolutionarily conserved. Taken together, our results reveal a unidirectional mechanical coupling between cell mechanics and nuclear mechanics via the regulation of Lamin A/C. We anticipate that mechanism of Lamin A/C regulation that we revealed, could form the basis for understanding nuclear mechanotransduction in epithelial tissues.

## INTRODUCTION

Mechanotransduction – a process through which mechanical forces are converted to biochemical signaling or gene expression is crucial for physiology^1^. Impaired mechanotransduction is at the heart of various diseases^2^. Mechanotransduction could be activated by either extrinsic forces like fluid shear flow in blood vessels^3^ or intrinsic forces through actomyosin contractility. Recent experiments have highlighted the importance of the transmission of forces to the nucleus through the cytoskeleton^4,5,6,7^. In this process, the nuclear scaffold plays an important role in activating mechanotransduction in the nucleus^8^.

Nuclear Lamins are the primary component of the nuclear scaffold. Lamins are type V intermediate filaments^9^. They comprise of A-type and B-type Lamins. In vertebrates, A-type Lamins are composed of two splice isoforms of the Lamin A gene, Lamin A and Lamin C, whereas B-Type Lamins are composed of Lamin B1 and Lamin B2. In contrast, in invertebrates like Drosophila, A-Type Lamin are composed of Lamin C and B-type Lamin is composed of Lamin DM_0_ ^10^. Lamins are known to influence mechanical properties of the nucleus. Lamin A/C contributes to the stiffness and viscosity of the nucleus whereas Lamin B is responsible for the elasticity of the nucleus. Recent experiments have shown that concentration of Lamin A/C is crucial for mechanotransduction in the nucleus. Not only does Lamin influence force induced stiffening of the nucleus^11^, it also promotes, nuclear translocation of MKL, a co-factor of Serum Response factor (SRF)^12^. Interestingly, Lamin B is expressed uniformly in all tissues during development, but Lamin A/C is developmentally regulated and has tissue specific expression profiles^13^.

Over the last decade, studies have focused on identifying how Lamin A/C is regulated in different tissues. Experiments in mesenchymal stem cells (MSC) and mesenchymal tissues have shown that Lamin A/C scales with ECM stiffness in these cells^14^. Epithelial tissue – a differentiated form of MSC, is one of the most abundant adult tissue. ECM in these tissues is scant ^15^, but the cells adhere to each other through a plethora of cell-cell junctions along the apico-basal axis^16^. These cell-cell junctions bear most of the mechanical stress in the tissue^17^. In the absence of strong interactions with the ECM, it is still unclear how Lamin A/C levels are regulated in these tissues. Studying this would be key to understand the role of Lamin A/C in mechanotransduction and the interplay between tissue mechanics and nuclear mechanotransduction in epithelial tissues.

In this work, using *Drosophila* epithelial tissues and mammalian Madin Darby Canine Kidney (MDCK) cells as a model system, we reveal that apico-basal cell compression is an ECM independent mechanism for regulation of Lamin A/C in epithelial tissues. Nuclear deformation in response to apico-basal cell compression modulates Lamin A/C levels. By combining genetic perturbations in vivo, and altered cell packing, in mammalian cells in culture, we show that apico-basal cell compression induced regulation of Lamin A/C is an evolutionarily conserved mechanism in epithelial tissues.

## RESULTS

### Epithelial tissues have heterogenous levels of Lamin C

As a first step towards investigating how Lamin A/C is regulated in epithelial tissues, we measured the concentration of Lamin C in epithelial tissues. We dissected and immunostained Salivary gland^18^, trachea^19^ and wing disc^20^ epithelial tissues from late third instar *Drosophila* larvae (Fig 1a) using an antibody against Lamin C (LamC), the only isoform of Lamin A/C expressed in *Drosophila*. We estimated the concentration of nuclear LamC by measuring the mean pixel intensity from a maximum intensity projection of a z-stack image (Fig. 1l and Online Methods). We found that LamC levels are low in the nuclei of salivary gland (SG) cells (Fig 1 b-c) and highest in the nuclei of tracheal cells (Fig 1 f-k). We observed significant differences of LamC levels even within a single wing disc. When we measured the concentration of LamC in different regions of the wing disc we found that LamC is very low in the wing pouch (Fig 1 j-k). LamC is about 3-fold higher in the fold regions of the wing and about 4-fold higher in the PM cells (Fig 1 h-i). Upon comparing LamC concentration in the SG and trachea with the wing disc pouch, we observed that the concentration of LamC in the SG is similar to that of the wing pouch, whereas tracheal cells have about 5-fold higher concentrations of LamC (Fig. 1m). We asked whether such large differences are specific to LamC by immunostaining Lamin DM_0_, ortholog of Lamin B in *Drosophila* and is known to be expressed throughout development^13^. Interestingly, we found no significant differences in the concentrations of LamDM_0_ in *Drosophila* tissues (Supplementary Fig. 1). These results indicate that different epithelial tissues have different levels of LamC and that the observed differences are not an artefact of immunostaining or deep tissue imaging.

**Figure 1:**
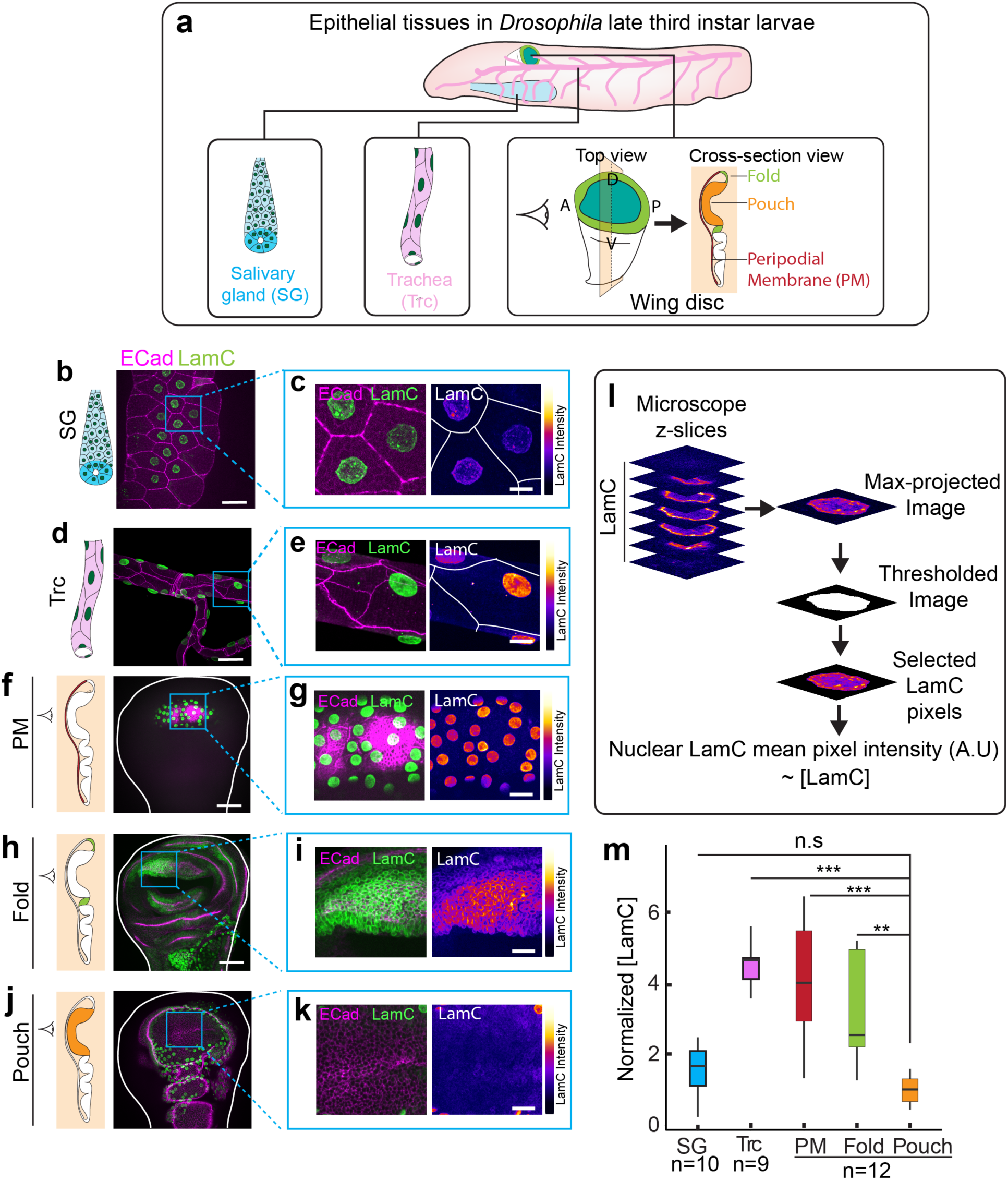
LamC is heterogeneously expressed in epithelial tissues. **(a)** Schematic showing different epithelial tissues in Drosophila. **(b)** LamC in salivary glands. Image of trachea showing E-Cad (magenta) and LamC (green). **(c)** Enlarged image of the region marked by the blue ROI in (b). Left panel shows the merge of E-Cad and LamC. Right panel shows the color coded LamC intensity. Solid white lines represent cell boundaries. **(d)** LamC in trachea. Image of salivary gland showing E-Cad (magenta) and LamC (green). **(e)** Enlarged image of the region marked by the blue ROI in (e). Left panel shows the merge of E-Cad and LamC. Right panel shows the color coded LamC intensity. Solid white lines represent cell boundaries. **(f)** LamC in trachea. Image of PM showing E-Cad (magenta) and LamC (green).**(g)** Enlarged image of the region marked by the blue ROI in (f). Left panel shows the merge of E-Cad and LamC. Right panel shows the color coded LamC intensity. **(h)** LamC in the fold. Image of fold showing E-Cad (magenta) and LamC (green). **(i)** Enlarged image of the region marked by the blue ROI in (h). Left panel shows the merge of E-Cad and LamC. Right panel shows the color coded LamC intensity. **(j)** LamC in the pouch. Image of pouch showing E-Cad (magenta) and LamC (green). **(k)** Enlarged image of the region marked by the blue ROI in (j). Left panel shows the merge of E-Cad and LamC. Right panel shows the color coded LamC intensity. **(l)** Schematic showing the estimation of LamC concentration in the nucleus of epithelial tissues. **(m)** Box-plot showing the normalized concentration of LamC in the epithelial tissues. Normalization is performed with respect to the concentration in the pouch. The sample number, n represents the number of tissue samples. One-way ANOVA was performed to estimate p-values. The p-values are shown in comparison to LamC concentration in the pouch. *** represents p< 10^−6^, ** represents p< 0.001 and n.s represents the differences are not significant. Scale bar in overview images (b,d,f,h,j), 25 µm. Scale bar in enlarged images, 15 µm.

### Lamin C scales with apico-basal cell compression in epithelial tissues

Next, we investigated the origin of the differences in LamC concentration across *Drosophila* epithelial tissues. A characteristic feature of epithelial tissues is their cell packing density, defined as the number of cells per mm^2^ surface area of the tissue. Squamous epithelial tissues have a very low packing density, whereas pseudostratified tissues have a very high packing density. As a consequence of the low packing density in squamous tissues, cells are deformed apico-basally. We examined whether LamC could scale with the cell packing density or apico-basal deformation of cells. To do so, we measured the apical cell area (*A*) and apico-basal height (*h*) in SG, trachea and different regions of the wing disc (Fig 2 a-e). In order to measure apico-basal deformation of cells, we approximated the cells in the epithelia as cuboids and defined an apico-basal cell deformation index (Fig 2f) as,

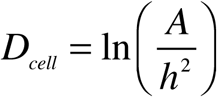

**Figure 2:**
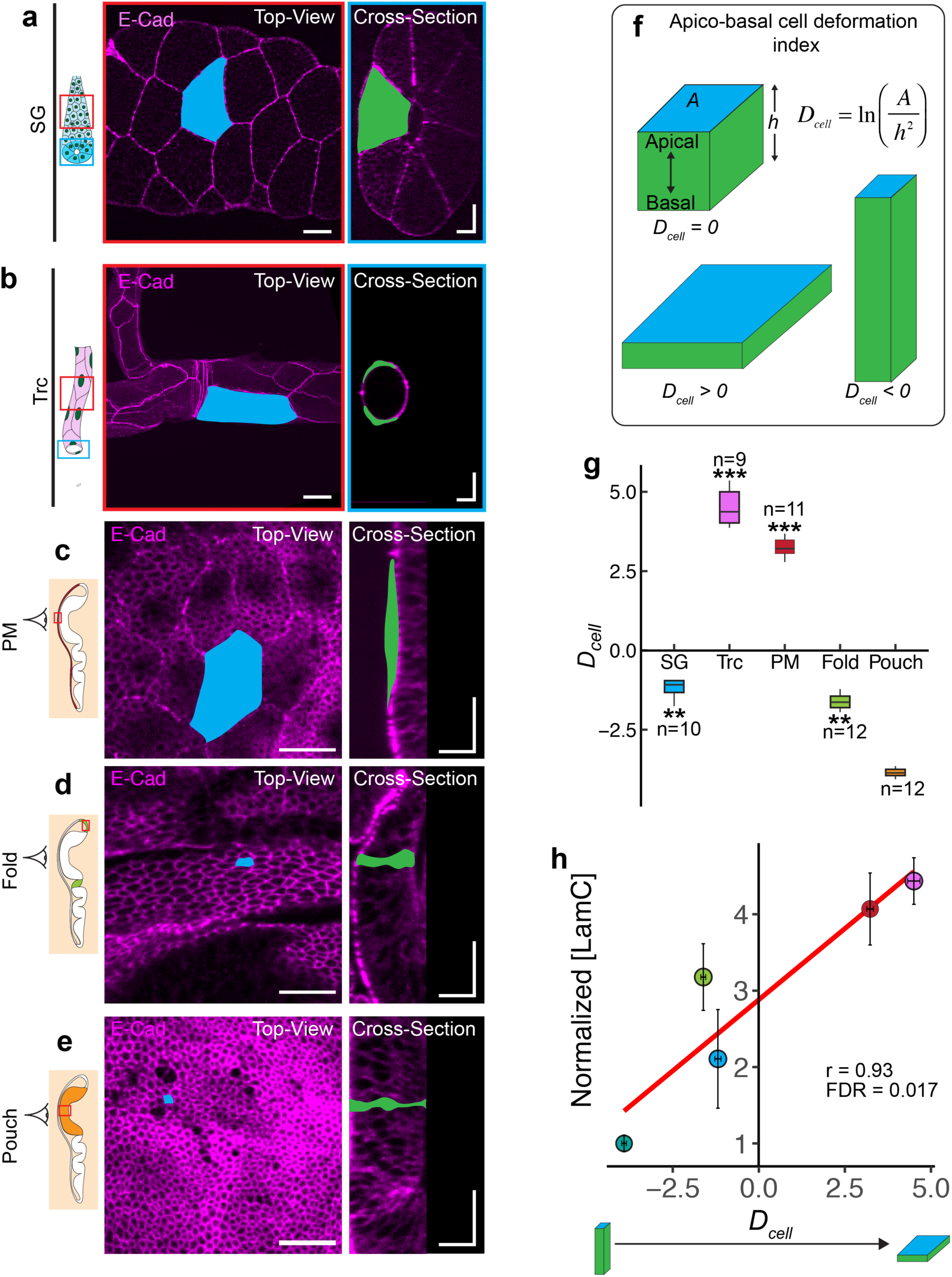
Lamin C scales with apico-basal cell compression in epithelial tissues. **(a-b)** Schematic representation of the salivary gland (a) and trachea (b) with the region imaged marked by the red and blue ROIs. Images show the top view (red ROI) and cross-section view (blue ROI) of the SG and trachea expressing E-Cad^GFP^ (magenta). A representative cell is shown with apical area marked in blue and apico-basal area marked in green. Scale bar in top-view images in (a-b), 25 µm. Scalebar in cross-section images, 25 µm along both axes.(**c-e**) Cell morphology in different regions of the wing disc. PM of the wing disc is colored in dark red (c). The fold is colored in light green (d), and the pouch is colored in orange (e). Images show the top view and cross-section view of each wing disc region expressing E-Cad^GFP^ (magenta). A representative cell is shown with the apical area marked in blue and apico-basal area marked in green. Scale bar in top-view images, 15 µm. Scale bar in cross-section images, 15 µm along both axes. **(f)** Schematic representation of cell deformation index (*D*_*cell*_) for epithelial tissues. *D*_*cell*_ for a cube is 1, whereas *D*_*cell*_ for a cuboid elongated along the apico-basal axis is < 1 and a cuboid compressed along the apico-basal axis is > 1. **(g)** Box plot of *D*_*cell*_ for different epithelial tissues of *Drosophila*. The sample number, n represents the number of tissue samples **(h)** Scatter plot between *D*_*cell*_ and normalized concentration of LamC. Each point in the scatter plot represents a tissue in *Drosophila* and the data is represented as mean± S.E.M. Solid red line is the linear regression to the scatter plot. The Pearson correlation coefficient is 0.93 and the false discovery rate (FDR) of the correlation is 0.017. P-values are estimated by one-way ANOVA and the p-values are shown for comparison with the LamC concentration in the pouch. ** represents p < 0.001 and *** represents p < 10^−5^.

We defined *D*_*cell*_ such that it is close to 0 for cuboidal cells, negative for columnar cells that are elongated along the apico-basal axis, and positive for squamous cells, compressed along the apico-basal axis. In accordance with previous studies^21^, we found that for the SG, both apical area and apico-basal height are very large, resulting in a *D*_*cell*_ close to 1. In contrast, both the trachea and PM have large apical areas but very small apico-basal heights, leading to large values of *D*_*cell*_ (Fig. 2g and Supplementary Fig. 2), as shown previously^22,23^. In the wing disc pouch, the apical area is small, but the apico-basal height is large, resulting in a small *D*_*cell*_. The *D*_*cell*_ values in these different tissues of *Drosophila* vary across four orders of magnitude. In order to observe the influence of *D*_*cell*_ on LamC levels in *Drosophila* tissues, we plotted the concentration of LamC in these tissues against the *D*_*cell*_ values (Fig 2h). We found that LamC levels showed a linear correlation with *D*_*cell*_, with a Pearson’s correlation of 0.93. The false discovery rate (FDR) for the correlation was 0.017 indicating a strong correlation. These results show that LamC in epithelial tissue scales with the apico-basal compression of cells.

### LamC is not regulated by ECM concentration in epithelial tissues

Since LamC levels in mesenchymal tissues are known to scale with ECM concentration^14,24^, we investigated whether such scaling exists also for epithelial tissues. Towards this end, we visualized collagen IV in *Drosophila* tissues (Fig 3a). We estimated the concentration of Collagen IV in the trachea, salivary gland and wing disc by the measuring mean fluorescence intensity (intensity/pixel) of an endogenously tagged Collagen IV (vkg^GFP^) from maximum intensity projected images. We found that the concentration of collagen IV is similar in the Salivary gland and trachea, but the levels of Lamin C are significantly higher in the trachea. In the wing disc, the Collagen IV concentration was found to be similar in the squamous PM and cuboidal cells of the disc fold but significantly higher in the wing disc pouch (Fig 3b). In contrast, Lamin C concentration is very low in the pouch as compared to the folds and the PM. Moreover, the observed difference in the Collagen IV concentration across the wing disc is not unique for this ECM protein. Other ECM proteins like Laminin-A (LanA), Laminin-B1 (LanB1) and Perlecan (Trol) also show similar concentration profiles in the wing disc (Supplementary Fig. 3a-f). Interestingly, when we plotted the concentration of Lamin C against the concentration of Collagen IV across different epithelial tissues, we did not find any significant correlation between them (Fig. 3c) (Pearson Correlation coefficient, −0.57 with FDR of 0.18). These experiments show that unlike mesenchymal tissues, LamC levels in epithelial tissues do not scale with ECM concentration.

**Figure 3:**
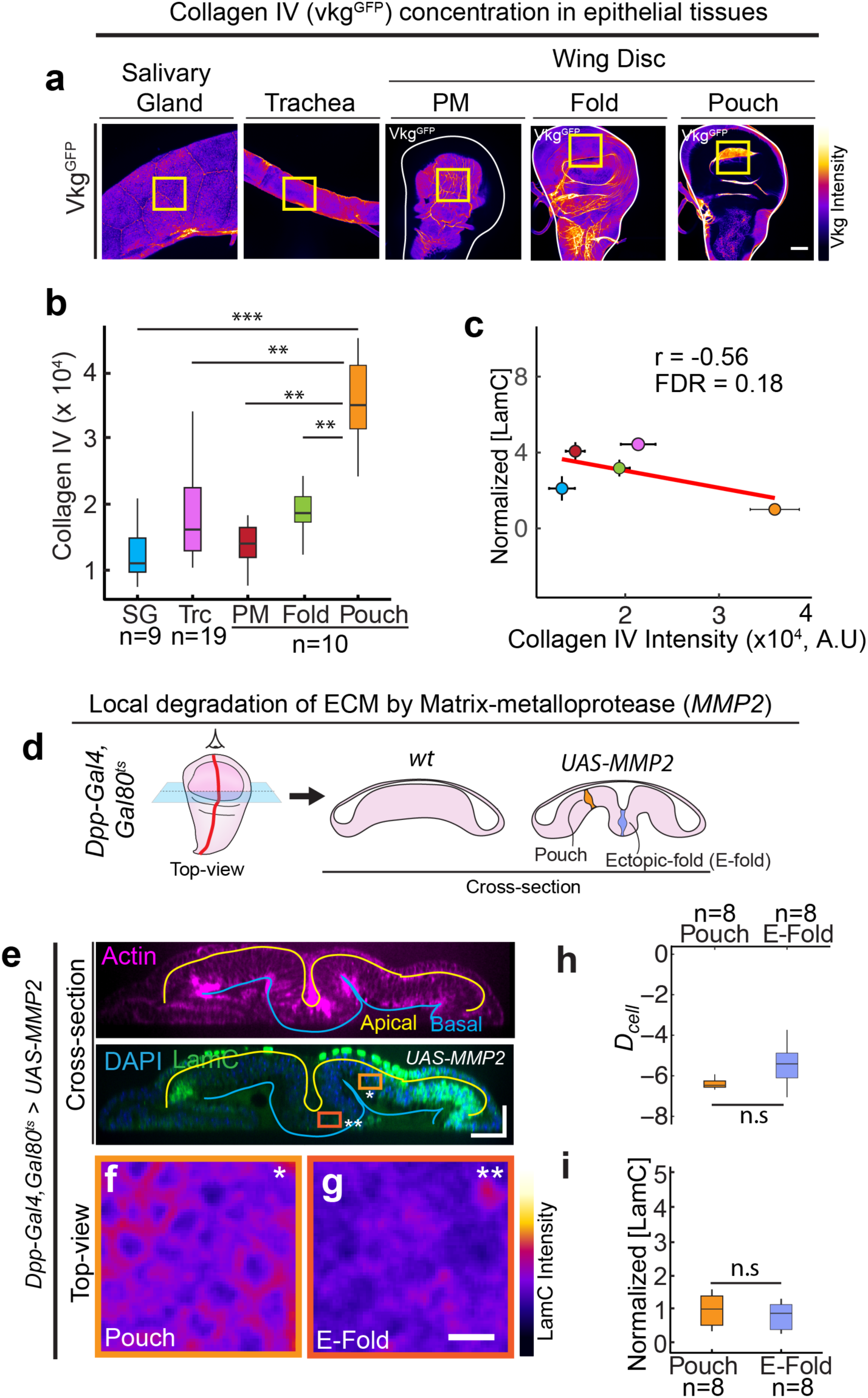
LamC is not regulated by ECM concentration in epithelial tissues. **(a)** Images showing Collagen-IV (encoded by Vkg in *Drosophila*) in different epithelial tissues. Yellow ROI represents the region of the tissue used to estimate the concentration of Collagen-IV. **(b)** Box plot shows the concentration of Collagen-IV in different epithelial tissues. **(c)** Scatter plot between Collagen IV and normalized LamC concentration in different tissues. Each point represents an epithelial tissue and the solid red line represents the linear regression to the points. The data is represented as mean ±S.E.M. The pearson correlation coefficient is −0.57 and the false discovery rate (FDR) of the correlation is 0.18. **(d)** Schematic showing the local degradation of ECM by overexpression of *MMP2* using Dpp-Gal4. The left panel shows the top-view of the wing disc and the right panel shows the cross section view of the wt wing disc and wing disc expressing *UAS-MMP2*. **(e)** Cross-section through the DV boundary of the wing disc expressing *UAS-MMP2* by *Dpp-Gal4*. The upper panel shows actin (magenta), and the lower panel shows the merge between LamC and DAPI. Solid yellow line shows the apical surface of cells, and the blue line shows the basal surface. Scale bar along both axes, 25 µm. **(f-g)** Color coded LamC images of the regions shown in (b). The pouch region is shown by the light orange ROI and single asterisk, and the ectopic fold (E-fold) region is shown by the dark orange ROI and double asterisk. Scale bar, 5 µm. **(h)** Box plot showing *D*_*cell*_ in wing pouch and E-fold. **(i)** Box plot showing normalized LamC in the pouch and the E-fold. The sample number, n represents the number of wing disc. Normalization is done with respect to the pouch. The p-values between two samples are estimated with a Student’s t-test and between multiple samples by one-way ANOVA. The p-values in (b) are shown in comparison to the concentration of collagen IV in the wing pouch. *** represents p < 10^−6^, **represents p < 10^−5^ and n.s represents that the differences are not significant.

Next, we investigated whether ECM is required for maintaining the LamC levels in the wing disc. To this end, we used Matrix metalloprotease to degrade the ECM^25^. We locally degraded the ECM in the wing disc by expressing *Drosophila* matrix metalloprotease (MMP2) in a spatially confined stripe using *Dpp-GAL4* (Fig. 3d-e). Degrading the ECM by expressing *UAS-MMP2* in the Dpp stripe is known to induce an ectopic fold in the wing disc^26^. We degraded the ECM for 24 hours in the late L3 stage of larval development by using a temperature sensitive Gal80 (Gal80^ts^). Temporal control is crucial, as prolonged ECM degradation is lethal for the larva. Upon degradation of ECM (visualized by Collagen IV) in the ectopic fold (Supplementary fig 3g), we observed a small decrease in cell height, in accordance with previous studies^26,27^ (Supplementary Fig. 4a). The deformation index (*D*_*cell*_), however, remained unchanged between *wt* and *UAS*-*MMP2* overexpression (Fig. 3h). Interestingly, LamC levels were not significantly different between pouch and ectopic fold (Fig 3f-g & Fig. 3i), consistent with our observation that LamC levels scales with *D*_*cell*_. Though previous experiments have shown that interaction of the cells with ECM is necessary to maintain cell shape in the wing disc^27^, our experiments clearly show that LamC levels in wing disc epithelia are not regulated by ECM concentration and thus also ECM stiffness.

### Apico-basal cell compression regulates LamC levels

To test whether the apico-basal compression of cells regulates LamC levels, we modulated cell shapes in the wing discs and observed its effect on LamC levels. First, we reduced the apico-basal height of the cells by overexpressing a dominant negative form of CDC42 (*CDC42*^*F89*^) in a stripe using *Dpp-GAL4* (Fig. 4a-b). CDC42 has been shown previously to regulate the apico-basal elongation of cells in the wing pouch^28^. Interestingly, overexpression of *CDC42*^*F89*^ induced a deep ectopic fold by reducing the height and apical cross-section of cells expressing *CDC42*^*F89*^ (Supplementary Fig. 4c-d). This perturbation resulted in a significant increase in *D*_*cell*_ of the cells in the ectopic fold as compared to the pouch region outside the fold (Fig. 4e). When we compared the LamC levels in the ectopic fold and the wing pouch outside of the fold, we found a significant increase in LamC in the ectopic fold (Fig. 4c-d & Fig. 4f). Next, we increased the height of the PM cells, which are usually very flat. We ectopically overexpressed a transcription factor Lines (*Lin*) in the PM using Ubx-Gal4 (Fig. 4g). Usually, *Lin* is expressed in the pouch but is absent in the PM. As shown previously^29,30^, we observed a significant increase in cell height upon overexpressing *Lin* in the PM (Fig. 4h & Supplementary Fig. 4e-f). We also observed that the cells overexpressing *Lin* have a reduced apical area as compared to *wt* (Supplementary fig. 4). When we measured *D*_*cell*_ for wt and *Lin* overexpression, we observed that in wt, *D*_*cell*_ is significantly higher in the PM as compared to the pouch (Supplementary Fig 4g). In contrast, upon *Lin* overexpression, *D*_*cell*_ in the PM and pouch are indistinguishable (Supplementary Fig 4g). We found that in the *Lin* overexpressed disc, the levels of LamC between the pouch and the PM are indistinguishable (Supplementary Fig. 4h). Interestingly, when we plotted the cell deformation index, *D*_*cell*_ and normalized LamC for all the perturbations, all these data followed the same linear correlation shown in Fig 2h (Fig. 4l), suggesting that the levels of Lamin A/C can be predicted solely from the apico-basal deformation of cells. These results clearly show that the apico-basal compression of cells regulates Lamin C levels in the wing disc.

**Figure 4:**
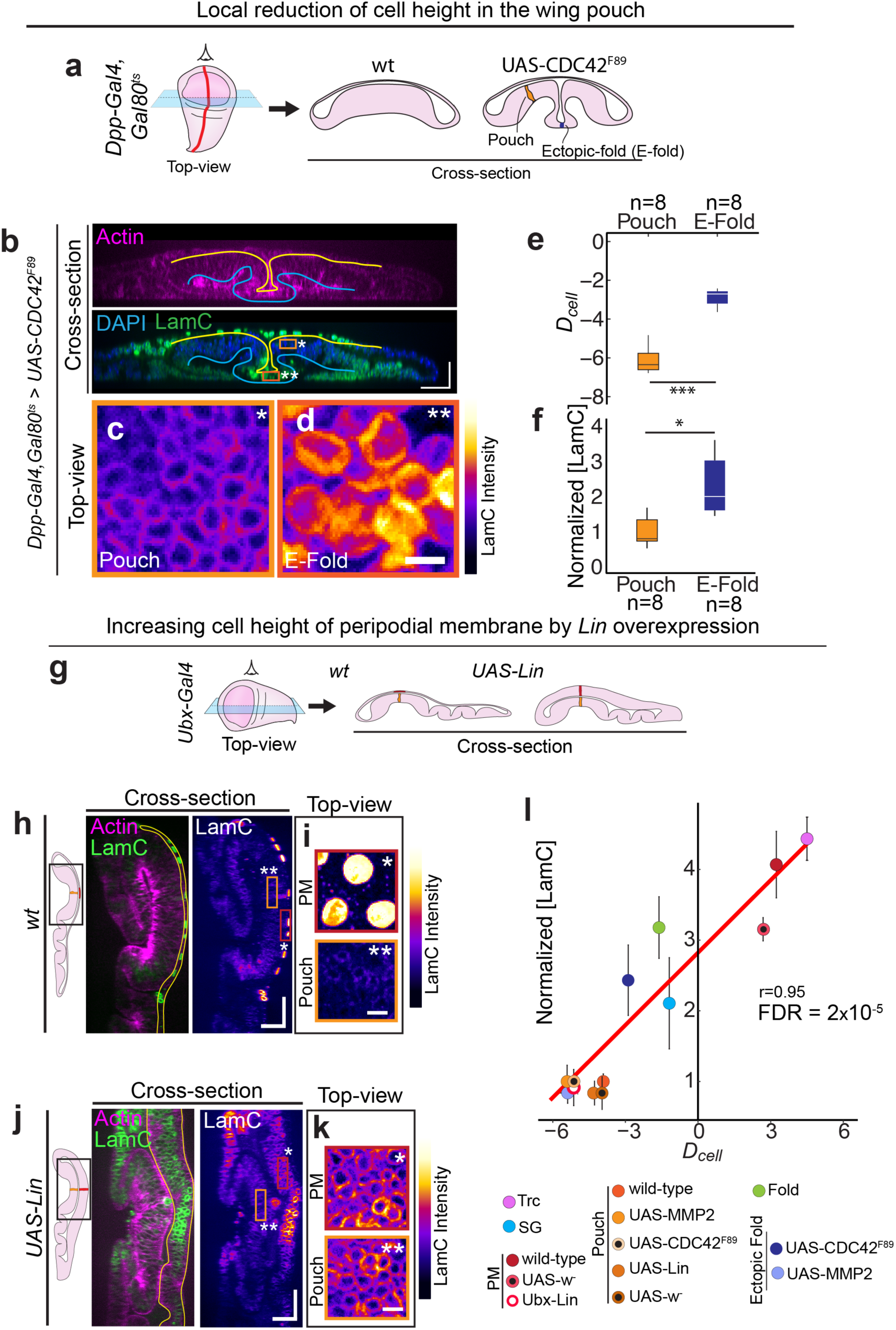
Apico-basal cell compression regulates LamC in epithelial tissues. **(a)** Schematic showing the local reduction in cell height by overexpression of a dominant negative form of *CDC42* using Dpp-Gal4. Left panel shows the top-view of the wing disc and the right panel shows the cross-section view of the wing disc. **(b)** Cross-section through the DV boundary of the wing disc expressing *CDC42*^*F89*^ by Dpp-Gal4 along the AP boundary. Upper panel shows actin (magenta) and lower panel shows merge between LamC and DAPI. Solid yellow line shows the apical surface, and the blue line shows the basal surface of the cells. Scale bar along both axes, 25 µm. **(c-d)** Color coded LamC images of the regions shown in (b). The pouch region is shown by the light orange ROI and single asterisk, and the ectopic fold (E-fold) region is shown by the dark-orange ROI and double asterisk. Scale bar, 5 µm. **(e)** Box plot showing *D*_*cell*_ in the wing pouch and E-fold. **(f)** Box plot showing normalized LamC in pouch and the E-fold. Normalization is done with respect to pouch. The sample number, n represents the number of wing discs. **(g)** Schematic showing the increase in cell height of PM cells by ectopic overexpression of *UAS-Lin* using Ubx-Gal4. Left panel shows the top-view of the wing disc and the right panel shows the cross-section view of the wing disc. **(h)** Cross-section of the *wt* wing disc in the region of the wing disc shown by the black ROI. Left panel shows merge of actin (magenta) and Lam C (green). Solid yellow line shows the PM. Right panel shows the color-coded image of LamC. Scale bar along both axes, 25 µm. **(i)** Top-view color-coded LamC intensity of the PM (marked by the dark-red ROI and single asterisk) and pouch (light-orange ROI and double asterisk). Scale bar, 5 µm **(j)** Cross-section images of wing disc with ectopic overexpression of *UAS-Lin* in the region shown by black ROI. Left panel shows merge of actin (magenta) and Lam C (green). Solid yellow line shows the PM. Right panel shows the color-coded image of LamC. Scale bar along both axes, 25 µm. **(k)** Top-view color-coded LamC intensity of the PM (marked by the dark-red ROI and single asterisk) and pouch (light-orange ROI and double asterisk). Scale bar, 5 µm (l) Scatter plot between Normalized LamC levels and *D*_*cell*_. The data includes different perturbations which change *D*_*cell*_ in the wing disc. Normalization is done with respect to the *wt* control in the respective experiment. The Pearson Correlation coefficient is 0.95 with FDR of 2×10^−5^. The p-values are estimated using a Student’s t-test. *** represents p < 0.001, * represents p< 0.05 and n.s indicates that the differences are not significant.

Next, we asked whether changes in LamC expression could also influence the morphology of epithelial cells. We first downregulated LamC in the PM of the wing disc, by driving LamC RNAi using Ubx-Gal4, which is primarily expressed in the PM. We found that in wing discs expressing LamC RNAi, LamC levels are significantly reduced as compared to wild-type (Supplementary fig. 5a & c). But, the apical cell area and height between wild-type and LamC RNAi knockdown cells are indistinguishable (Supplementary fig. 5b & d). Next, we overexpressed LamC in a stripe of cells using *Ptc-Gal4*. We restricted expression to the late L3 larval stage by using the temperature sensitive *Gal80*^*ts*^ and shifting the larvae from 25 °C to 29 °C (Supplementary fig. 5e). This perturbation causes significantly higher levels of LamC in cells along the stripe (Supplementary fig. 5e). However, the height of the cells in the stripe is identical to the cells outside the stripe (Supplementary fig. 5g-h). Even in LamC mutant larvae, the height of cells in the wing pouch is identical to the cells in the wild-type wing disc (Supplementary Fig 5i).These experiments clearly show that there is a unidirectional coupling between cell mechanics and LamC regulation in epithelial tissues - apico-basal compression of epithelial cells regulates LamC, but altering LamC does not influence cell morphology in epithelial tissues

### Apico-basal cell compression regulates Lamin A/C levels by flattening the nucleus

Next, we investigated whether apico-basal compression of cells induces nuclear deformation leading to increase in Lamin A/C levels. We visualized nuclear shape in Salivary glands, trachea and wing discs (Fig. 5 and Supplementary fig. 6a-b). We measured cross-sectional area (*A*_*n*_) and nuclear height (*h*_*n*_) in these nuclei. We defined the nuclear deformation index as,

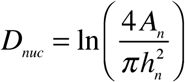

**Figure 5:**
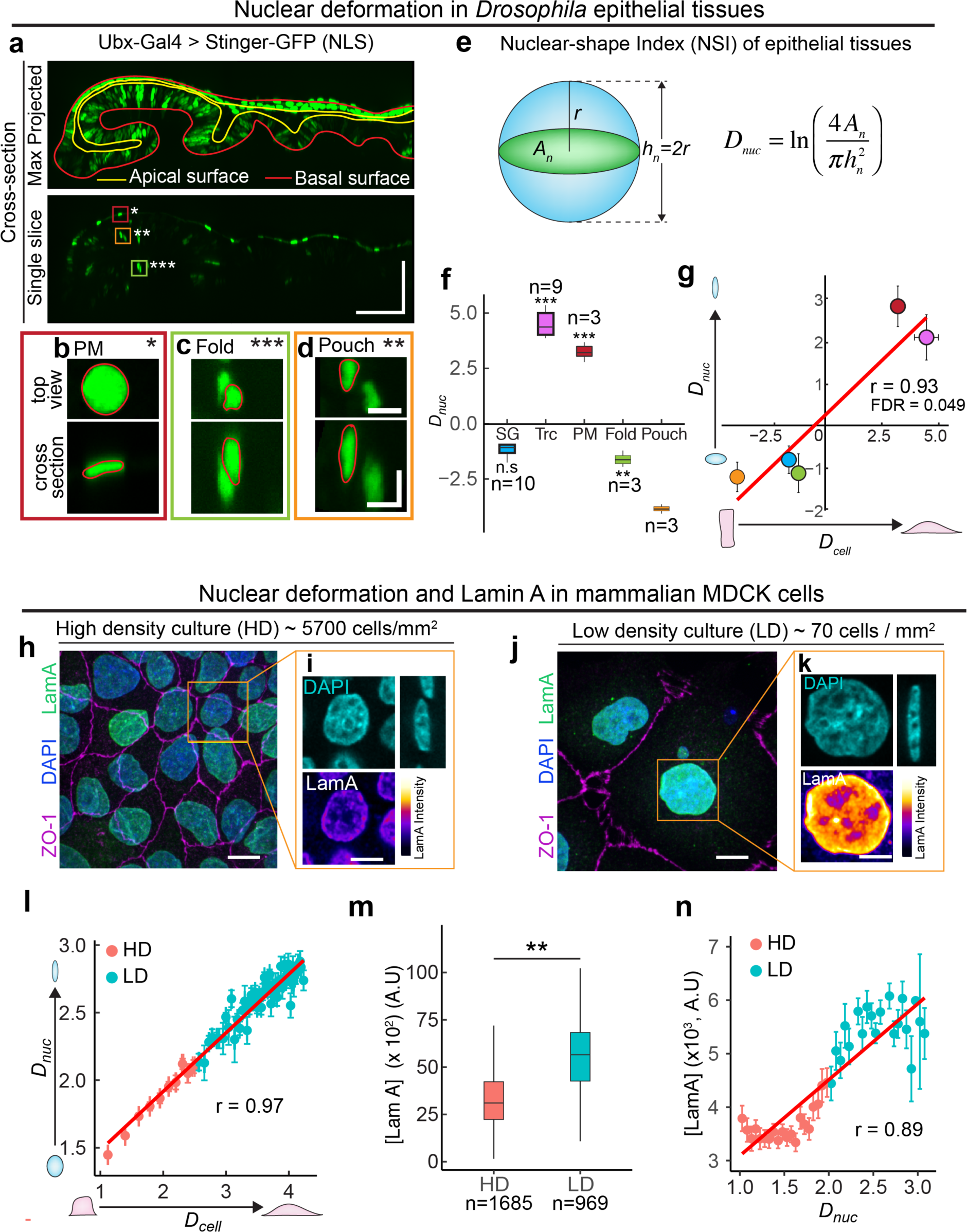
Apico-basal cell compression regulates Lamin A/C by flattening the nucleus. **(a)** Cross-section view of the wing disc expressing nuclear NLS protein Stinger-GFP driven by Ubx-Gal4. Upper panel shows the maximum projected image of Stinger-GFP. Solid red line shows the basal surface of cells, and solid yellow line shows the apical surface of cells. Scale bar along both axes, 50 µm. **(b-d)** Enlarged images of regions where PM is marked by dark-red ROI and single asterisk (b), fold by green ROI and triple asterisk (c) and pouch by orange ROI and double asterisk (d). For each region, the upper panel shows the top-view of the nucleus, and lower panel shows the cross-section of the nucleus. Solid red line shows the outline of the nucleus. Scale bar along both axes, 5 µm. **(e)** Schematic showing the estimation of nuclear deformation index (*D*_*nuc*_) for a nucleus. **(f)** Box plot showing *D*_*nuc*_ for different *Drosophila* tissues. The sample number, n represents the number of tissue samples **(g)** Scatter plot between *D*_*cell*_ and *D*_*nuc*_ in *Drosophila* tissues. Data is presented as mean± S.E.M. The Pearson Correlation coefficient is 0.93 with FDR of 0.049. **(h)** Image showing high density culture of 5700 cells/mm^2^. Tight junction protein ZO-1 is shown in magenta, Lamin A in green and DAPI in blue. Scalebar, 15 µm. **(i)** Enlarged images of the region shown by the orange ROI in (h). DAPI is shown in blue and Lamin A is shown in color coded intensity. Scale bar, 10 µm. **(j)** Image showing low density culture of 70 cells/ mm^2^. Tight junction protein ZO-1 is shown in magenta, Lamin A in green and DAPI in blue. Scalebar, 15 µm. **(k)** Enlarged images of the region shown by orange ROI in (j). DAPI is shown in blue, and Lamin A is shown in color-coded intensity. Scale bar, 10 µm. **(l)** Binned scatter plot between *D*_*cell*_ and *D*_*nuc*_ generated by binning *D*_*cell*_ in bins of 0.07. Each point in the plot represents mean value of *D*_*cell*_ and *D*_*nuc*_ in the bin. The red points represents data from HD culture and cyan points represents data from LD culture. The data are presented as mean± S.E.M. Solid red line represents the linear regression to the data. **(m)** Box plot showing LamA in high density (HD) and low density (LD) cultures. The sample number, n represents the number of cells analysed.**(m)** Binned scatter plot between *D*_*nuc*_ and LamA intensity generated by binning *D*_*nuc*_ in bins of 0.06. Each point in the plot represents mean value of *D*_*nuc*_ and LamA in the bin. The red points represents data from HD culture and cyan points represents data from LD culture. The data are presented as mean± S.E.M. Solid red line represents the linear regression to the data. The p-values are estimated by Student’s t-test, ** represents p < 0.01 and *** represents p < 10^−5^.

We defined *D*_*nuc*_ such that *D*_*nuc*_ = 0 for a spherical nucleus and deviates from 1 as the nuclear shape deviates away from a sphere. *D*_*nuc*_ is negative for prolate shapes and *D*_*nuc*_ is positive for oblate shapes. The nuclei of Salivary glands have a large cross-sectional area and nuclear height (Supplementary fig. 6a-b), making *D*_*nuc*_ close to 1. In contrast, trachea have a significantly larger *D*_*nuc*_, indicating that nuclei are much flatter in trachea as compared to salivary glands (Fig 5f). Measuring nuclear shape in the wing disc is challenging because segmentation of the densely packed nuclei in the disc pouch is extremely difficult. We overcame this problem by driving a nuclear localized GFP called “Stinger-GFP” with Ubx-Gal4 (Fig 5a). This Gal4 primarily expresses in the PM but has a patchy expression in the wing pouch (Supplementary Fig 6c). Thus, nuclei in the wing pouch and fold are sparsely labeled, enabling nuclear segmentation (Fig 5a). When we measured the cross-sectional area and height of the nuclei in the pouch and fold (Supplemntary Fig 6d-e), we found that the nuclei have a very low *D*_*nuc*_ and they are prolate (Fig 5f). The nuclei in the PM have very large *D*_*nuc*_ and they are oblate. Interestingly, *D*_*nuc*_ and *D*_*cell*_ in *Drosophila* tissues are strongly correlated, with a correlation coefficient of 0.93 (Fig 5g). These results show that the observed changes in LamC in *Drosophila* tissues is due to deformation of the nucleus in response to apico-basal cell deformation.

Next, we asked whether such apico-basal cell compression dependent LamC regulation is conserved in mammalian cells. To this end, we cultured mammalian Madin Darby Canine Kidney (MDCK) epithelial cells on a cover glass coated with Collagen I. In order to mimic the cell packing in the wing disc pouch and PM, we cultured the cells either at high density (∼5700 cells/mm^2^) or at low density (∼70 cells/mm^2^) (Fig 5h-j). We cultured both low density and high density cultures on the same culture dish coated with Collagen I to provide an ECM independent microenvironment. We observed that the apical area of the cells grown at high density were much smaller than that grown at low density (Fig 5 h-k). Concomitantly, the apico-basal height of cells grown at higher density was significantly larger than those grown at low density (Supplementary Fig 6h). We found that the nuclear deformation index, *D*_*nuc*_ is strongly correlated with cell deformation index, *D*_*cell*_, with a correlation coefficient of 0.96 (Fig. 5l). These results in *Drosophila* tissues and MDCK cells in culture show that nuclear deformation by the apico-basal cell compression is an evolutionary conserved phenomenon in epithelial tissues.

Next, we visualized Lamin A/C in MDCK cells by immunostaining the cells with an antibody against Lamin A (LamA) – a canine ortholog of *Drosophila* LamC. We performed the staining of both low-density and high-density cultures in the same culture dish, to minimize variation in the concentration of LamA in low density and high density cultures. We found that concentration of LamA was significantly higher in low density cultures than in high density cultures (Fig. 5m). The concentration and total amount of LamA in the nucleus strongly correlates with *D*_*nuc*_, with a Pearson’s correlation coefficient of 0.97 (Fig. 5n and Supplementary Fig. 6f). These results show that the regulation of Lamin A/C by apico-basal compression of cells is an evolutionarily conserved mechanism in epithelial tissues.

## Discussion

Lamin A/C is an important element of the cellular mechanotransduction pathway which bridges mechanical forces to the cell nucleus. In mesenchymal tissues Lamin A/C is known to be regulated by ECM stiffness^14^. How Lamin A/C is regulated in epithelial tissues was not known. In this work, we revealed a mechanism of Lamin A/C regulation in epithelial tissues by apico-basal cell compression. We found that in many different tissues and under different genetic perturbations Lamin A/C levels can be reliably predicted solely based on apico-basal cell compression. We also showed that in response to apical-basal cell compression nuclei deformed. Nuclear deformation is a common feature of Lamin A/C regulation in both mesenchymal^31^ and epithelial tissues. In both these tissues, Lamin A/C levels increase as nuclei change their shape from being prolate to oblate. In mesenchymal tissues cell nuclei deform by forces generated in actomyosin cables that link to the ECM^32,33^. In contrast, epithelial tissues alter cell shape by modulating cortical tension and properties of cell-cell junctions^28,34^. We show here that the resulting apico-basal compression can induce nuclear flattening and regulate Lamin A/C levels.

Interestingly, we also found that while cell morphology regulates Lamin A/C levels, the reverse is not true: changes in Lamin A/C levels do not influence the morphology of cells in *Drosophila* epithelial tissues. Even in Lamin C null mutants the morphology of the cells in the wing disc are not significantly different from wild-type, suggesting that altering Lamin A/C levels does not affect cytoskeletal organization in these cells. However, as previously described, the Lamin C null larvae do not survive beyond the late third instar stage of development^35,36^. This finding is in accordance with the role of Lamin A/C in protecting the genome against DNA damage^37,38^ and in mediating nuclear mechanotransduction^39^. Thus, in LamC null larvae, hindered translocation of mechanotransducers like YAP^40^ and MKL^41^ or accumulation of DNA damage during larval stages could lead to lethality at later development.

Taken together, our results suggest that regulation of Lamin A/C could be the underlying mechanism coupling cell mechanics and nuclear mechanotransduction. The nature of this mechanism and how nuclear mechanotransduction could influence developmental process are major questions that our study raises for future investigations.

## Acknowledgements

We are grateful for Christian Dahmann for providing the *CD8-Cherry;Dpp-Gal4,Gal80*^*ts*^ fly line. We are grateful to Lori Wallrath, for the UAS-LamC and LamC^EX296^/CyO fly lines. We thank the light microscopy facility at the MPI-CBG for help with the imaging. We thank Valentina Greco for valuable discussions and critical reading of the manuscript. We also thank Christian Dahmann, Shovamayee Maharana and Suhrid Ghosh for critical review of the manuscript prior to submission. This work was supported by funding from the Deutsche Forschungsgemeinschaft: EA4/10-1, EA4/10-2 (N.A.D., K.V.I, S.E.). Lastly, we dedicate the manuscript to the memory of our co-author Suzanne Eaton, who tragically passed away during the conceptualization of this work.

## Author Contributions

The project was conceived by K.V.I. Experiments were performed by K.V.I. Analysis was performed by K.V.I. The first draft of the paper was written by K.V.I with subsequent contributions by N.A.D and F.J.

## Online Methods

### Fly Genetics

All flies were kept at normal cornmeal food and maintained at 25 °C unless otherwise specified. The fly stock, w-;DECAD^GFP^ fly was a kind gift from the Hong lab (Huang et al). The fly stock, *w*^*-*^;*UAS-CD8::Cherry;Dpp-GAL4,Gal80*^*ts*^ was a kind gift from Christian Dahmann. The fly stocks *yw;vkg::GFP* and *w*^*-*^;*Ptc::Gal4,Gal80*^*ts*^ were obtained from the fly facility of Max Planck Institute of Molecular Cell Biology and Genetics, Dresden. The following fly stocks were obtained from Bloomington Drosophila Stock Center: *w*^*-*^;*UAS-mmp2* (#58705),, *w*^*-*^;*UAS-Lin (#7074), w*^*-*^;*UAS-CDC42*^*F89*^ (#6286*), w*^*-*^;*trol::GFP* (#60214*), w*^*-*^;*UAS-Stinger::GFP, w*^*-*^; *Df(2R)trix/CyO* (#1896), The following flystocks were obtained from Vienna Drosophila Stock Center: w-;Laminin-A::GFP (#318155), w-;Laminin-B1::GFP (#318180). The fly stocks *w*^*-*^;*UAS-LamC, w*^*-*^; *LamC*^*EX296*^*/CyO* were a kind gift from Dr. Lori Wallrath, University of Iowa. The fly stock *w*^*-*^;*UAS-LamC*^*RNAi*^ was obtained from Fly stocks of National Institute of Genetics (#10119R-1). The fly stocks, *w-;UAS-MMP2,Vkg::GFP* and *w-;CD8::Cherry; Ubx-GAL4* were generated in this work.

### Temperature Shift experiments

Temperature shift was used to activate flies expressing *Gal80*^*ts*^. In the experiment, where *Dpp-Gal4* is expressed under the control of *Gal80*^*ts*^, the larvae were grown at 18 °C till late third instar stage. As soon as first larvae come out of the food and start upcrawling (about 10-11 days at 18°C), the vial is shifted from 18°C to 29°C to activate the *Gal80*^*ts*^. The vial is kept at 29 °C for 24 hours before dissecting and fixing the wing disc.

### Dissection of Drosophila tissues

Late third instar larvae at upcrawling stage were used for all experiments. The dissecting protocols for different *Drosophila* tissues are given below:

#### Wing disc

Larvae were selected and transferred to ice cold 1x PBS for 5 min till the larvae are immobilized. Then the larvae were held by a pair of no. 55 forceps at one-third the body length from the anterior end and then the posterior part of the larvae is ripped with the help of another pair of no. 55 forceps. Then the mouth of the anterior part of the larvae is held by one pair of forceps and the larva is turned inside out revealing the imaginal discs. All the other tissues are removed leaving the just the wing imaginal disc attached to the body wall.

#### Salivary gland

The larvae were held at the middle by one pair of forceps and the mouth is held by another pair of forceps. The mouth is slowly pulled. As soon as the mouth parts start to come out, the forceps holding the middle of the larva is slowed released. This leads to the ejection of salivary glands by the fluid pressure inside the larva. The Salivary gland is cleaned off from the fat bodies attached and then collected in ice cold 1X PBS for fixation.

#### Trachea

Larvae were help by the mouth and the posterior end of the larva was pinned into a PDMS pad. Then the anterior end is also pinned into the PDMS pad. Then the larva is ventrally dissected using fine forceps. Then the fat body, gut and other organs are removed. Then the body wall is pinned to the sides and the rest of unwanted body parts are removed leaving the trachea attached to body wall.

### Immunofluorescence staining of Drosophila tissues

Different tissues that are dissected are fixed in 4% PFA for 20 min. Then the tissues are washed twice with PBX2 (1X PBS + 0.05% Triton X-100), 15 min for each wash. Then the tissues are washed in BBX250 (PBX2 + 1mg/ml BSA + 5mM NaCl) for 45 min. This was followed by overnight incubation at 4 °C in primary antibody diluted 1:50 in BBX (PBX2 + 1mg/ml BSA). The following primary antibodies were obtained from Developmental studies hybridoma bank: mouse monoclonal antibody against Lamin C (Cat No. LC28.26), mouse monoclonal antibody against Lamin DM_0_ (Cat No. ADL67.10). The primary antibody was washed twice with BBX, 20 min for each wash followed by washing in BBX+4% normal goat serum once for 45 min. This was followed by incubation in secondary antibody either for 2-3 hours at room temperature or overnight at 4°C. The following secondary antibodies were obtained from Life Technologies: goat anti mouse Alexa-Fluo647 Flour, goat anti mouse Alexa-Fluo488. The secondary antibody was removed followed by washing twice with PBX2. Then the tissues were incubated in a mix of DAPI and either Alexa-Fluo488-Phalloidin, Alexa-Fluo568-Phalloidin or Alexa-Fluo660 Phalloidin. This was followed by washing twice with 1x PBS and the tissues are stored in PBS till the tissues are mounted. Before mounting, wing imaginal discs, salivary glands and trachea are dissected from the body wall. Slides are prepared for mounting by creating a thin channel by using two strips of double sided tape (Tesa). The tissue samples stored in 1x PBS are sucked in by a 100 µl pipette and transferred to the slide in between the two tape strips. The excess PBS is removed and the tissues are arranged. Then the sample is covered with 22×22 sized No.1 coverslip so that the coverslip sticks to the double-sided tape. Then Vectashield mounting medium (Vector laboratories) is added to one side of the coverslip and allowed to seep in by capillary effect. Once the entire sample in mounted, excess Vectashield is removed and the sides of the coverslip is sealed using transparent nail polish. The slides are then stored at 4 °C till they are imaged.

### Cell Culture

Madin Darby Canine Kidney (MDCK) cells were cultured in Minimum Essential Medium (MEM) (Life Technologies,), supplemented with non-essential amino acids (Life Technologies, Cat# 11140076) sodium pyruvate (Life Technologies,11360039) and 5% Fetal Bovine Serum (FBS) (Life Technologis). Cells are passaged using 0.25 % Trypsin-EDTA (Life Technologies). Cells were cultured on glass bottom dishes (Matek). A three-well cell culture insert (iBiDi, Cat # 80366) was inserted to form separate wells for low density and high-density culture, and the cover glass was treated with 100 µg/ml Collagen I PureCol solution (Advanced BioMatrix, Cat # 5005) and incubated for 2 hours. The dishes were rinsed once with 1x PBS before seeding cells on them. We seeded ∼ 5700 cells/mm^2^ for high density culture and 70 cells/mm^2^ for low density culture. The cells were cultured for 24-48 hours before fixation.

### Immunofluorescence staining of MDCK cells

Cell culture medium was removed and cells were washed twice with 1x PBS. Then the cells were fixed in 4% Paraformaldehyde for 20 minutes. This was followed by washing twice with 1X PBS and then incubated in 0.5% Triton X-100 for 20 minutes. Then the cells were blocked for 20 minutes in 1% BSA (made in 1X PBS). The cells were then incubated overnight with mouse monoclonal antibody against Lamin A (Thermo Fischer) and rabbit polyclonal antibody against ZO1, both diluted 1:500 in 1% BSA. Then primary antibody was removed and the cells were washed twice with 1X PBS followed by incubating in 1% BSA for 20 minutes. Then cells were incubated for 2-3 hours in Alexa 647 goat anti mouse and Alexa 488 goat anti-rabbit secondary antibody (Life technologies), both diluted 1:500 in 1% BSA.

### Imaging

#### Imaging of Drosophila tissues

Samples were imaged on an Olympus IX81 microscope equipped with a spinning disk module (Yokogawa) and back illuminated EMCCD. Different imaging parameters were used for imaging different samples. Confocal z-stack of Lamin C immunostained samples were acquired using 40x silicon immersion objective with 0.47 µm z-spacing. Images of fluorescently tagged ECM proteins were imaged using 30x silicon immersion objective with 0.5 µm z-spacing. Drosophila wing disc expressing Nuclear GFP (Stinger-GFP), driven by Ubx-Gal4 was imaged using a 60x silicon immersion objective with 0.18 µm z-spacing. All images were acquired using a back illuminated EMCCD camera with an exposure time of 100 ms and EM gain of 250.

#### Imaging of MDCK cells

MDCK cells were imaged on an Olympus IX81 microscope equipped with a spinning disk module (Yokogawa). Confocal z-stack of the cells were acquired using a 60x silicon immersion objective, 0.28 µm spaced apart on back illuminated EMCCD camera with an exposure time of 100 ms and EM gain of 250.

### Image Analysis

Analysis of images were performed using Fiji and MATLAB (Mathworks). We used separate analysis routines for analysis of images acquired for Drosophila tissues and MDCK cells.

#### Analysis of Drosophila tissues

Lamin C is present on the periphery of the cell nucleus. In order to analyse the concentration of nuclear Lamin, the maximum projection algorithm in FIJI is used to project the Lamin C intensity from different planes over the nucleus in a z stack. Then using a custom written program in MATLAB, the maximum projected plane is thresholded and multiplied with the maximum projected image to select pixels that belong to the nucleus. This is followed by calculating the mean intensity of Lamin C per pixel, which is a measure of concentration of Lamin C in the nucleus. Apical cell area and apico-basal height of the cells in Drosophila tissues were estimated using custom written programs in MATLAB

#### Analysis of MDCK cells

Cell morphology, nuclear morphology and Lamin C concentration in MDCK cells was estimated using custom written program in MATLAB. Fluorescence intensity of ZO-1 was used to estimate the apical cell area. ZO-1 network was segmented using Tissue Analyzer plugin in FIJI. The segmented image was inverted to estimate the apical cell area. Nucleus is segmented in 3-dimensions using MATLAB and the segmented nucleus is maximum projected along the z-axis and along the y-axis. The projection along the z-axis was used to estimate the cross-sectional area of the nucleus. An ellipse is fit to the projection of the nucleus along the y-axis and the minor axis of the ellipse is used as the height of the nucleus. The height of the cell is estimated from the height of the nucleus and the distance between the top of the nucleus and the top of the cell. In order to calculate the amount of Lamin A in the nucleus, the 3-D segmented nucleus is used as a mask and multiplied with the intensity image to select the pixels that belong to the nucleus. The total intensity if the masked 3-D nucleus is used as the total Lamin A intensity and the mean intensity per pixel is used as the concentration of Lamin A.

### Plotting and Statistical Analysis

All graphs presented in the paper are plotted using R-Studio and the figures are prepared using Adobe Illustrator. Statistical analysis is performed using MATLAB. One -way ANOVA is used when multiple quantities are compared, whereas Student’s t-test was used when two samples are compared. False discovery rate analysis is used to estimate the significance of correlations. One variable of the correlation is randomized iteratively for 105 iterations and Pearson Correlation Coefficient (PCC) is estimated for each iteration. Then then the fraction of iterations with PCC better than the original PCC is estimated as the false discovery rate. We consider FDR < 0.05 as a significant correlation.

## Supplementary Figures

**Supplementary Fig. 1:**
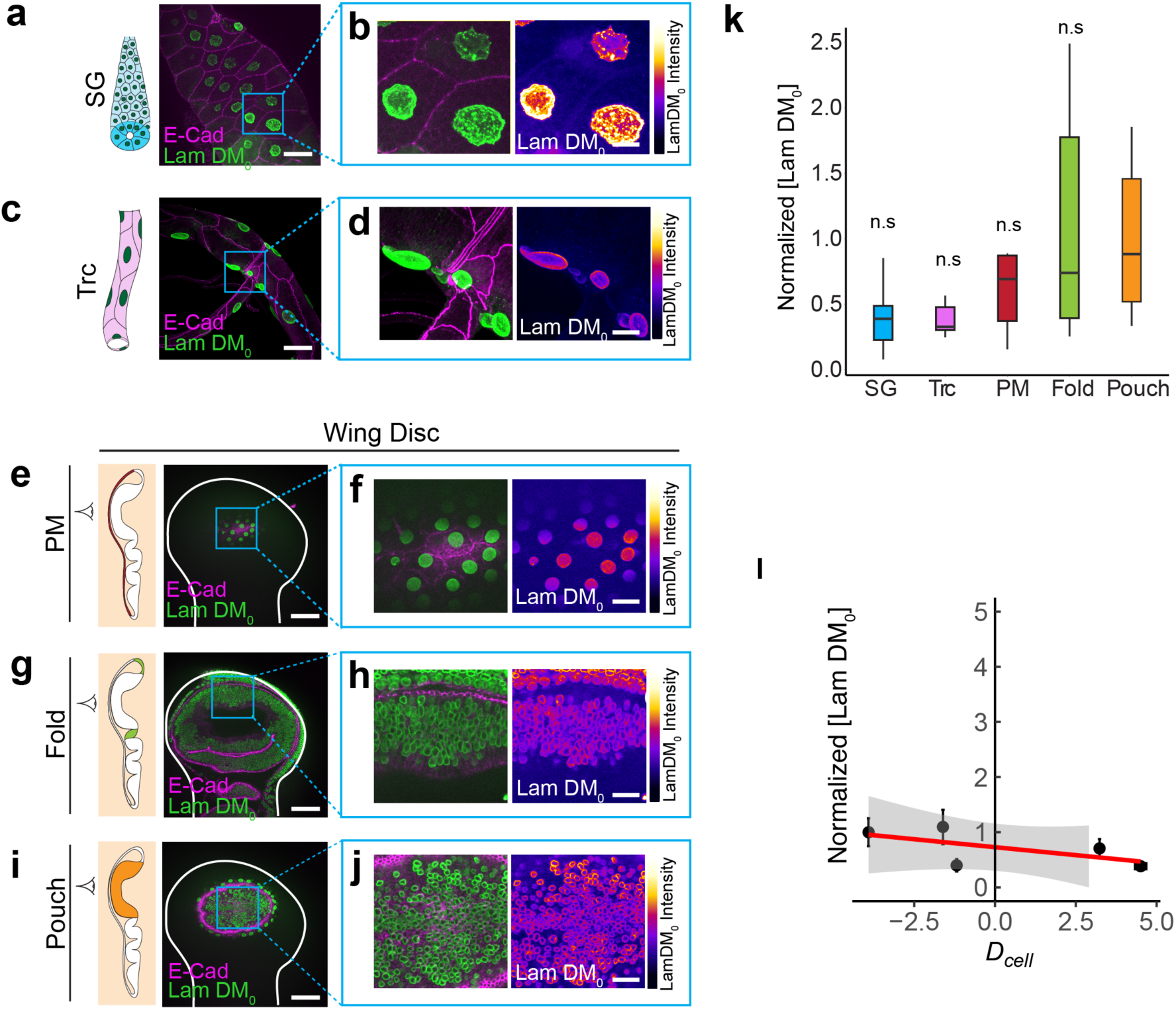
Lamin DM_0_ is similarly expressed in different epithelial tissues. **(a)** Image showing E-Cad (magenta) and LamDM_0_ (green) in salivary gland. **(b)** Enlarged image of the region marked by blue ROI in (a). Left panel shows the merge of E-Cad and LamDM_0_. Right panel shows LamDM_0_ color coded for intensity. **(c)** Image showing E-Cad (magenta) and LamDM_0_ (green) in trachea. **(d)** Enlarged image of the region marked by blue ROI in (c). Left panel shows the merge of E-Cad and Lam DM_0_. Right panel shows LamDM_0_ color coded for intensity. **(e)** Image showing E-Cad (magenta) and LamDM_0_ (green) in wing disc PM. **(f)** Enlarged image of the region marked by blue ROI in (e). Left panel shows the merge of E-Cad and LamDM_0_. Right panel shows LamDM_0_ color coded for intensity. **(g)** Image showing E-Cad (magenta) and LamDM_0_ (green) in wing disc fold. **(h)** Enlarged image of the region marked by blue ROI in (g). Left panel shows the merge of E-Cad and LamDM_0_. Right panel shows LamDM_0_ color coded for intensity. **(i)** Image showing E-Cad (magenta) and LamDM_0_ (green) in wing disc pouch. **(j))** Enlarged image of the region marked by blue ROI in (i). Left panel shows the merge of E-Cad and LamDM_0_. Right panel shows LamDM_0_ color coded for intensity. **(k)** Box plot showing the normalized LamDM_0_ concentration in different *Drosophila* tissues. Normalization is performed with respect to wing disc pouch. P-values were estimated by one-way ANOVA. Comparison is shown with wing pouch and n.s represents that the differences are not significant. **(l)** Scatter plot between *D*_*cell*_ and normalized LamDM_0_. Error bars indicate mean ± SE. Scale bar in overview images, 50 µm. Scale bar in enlarged images, 15 µm.

**Supplementary Fig. 2:**
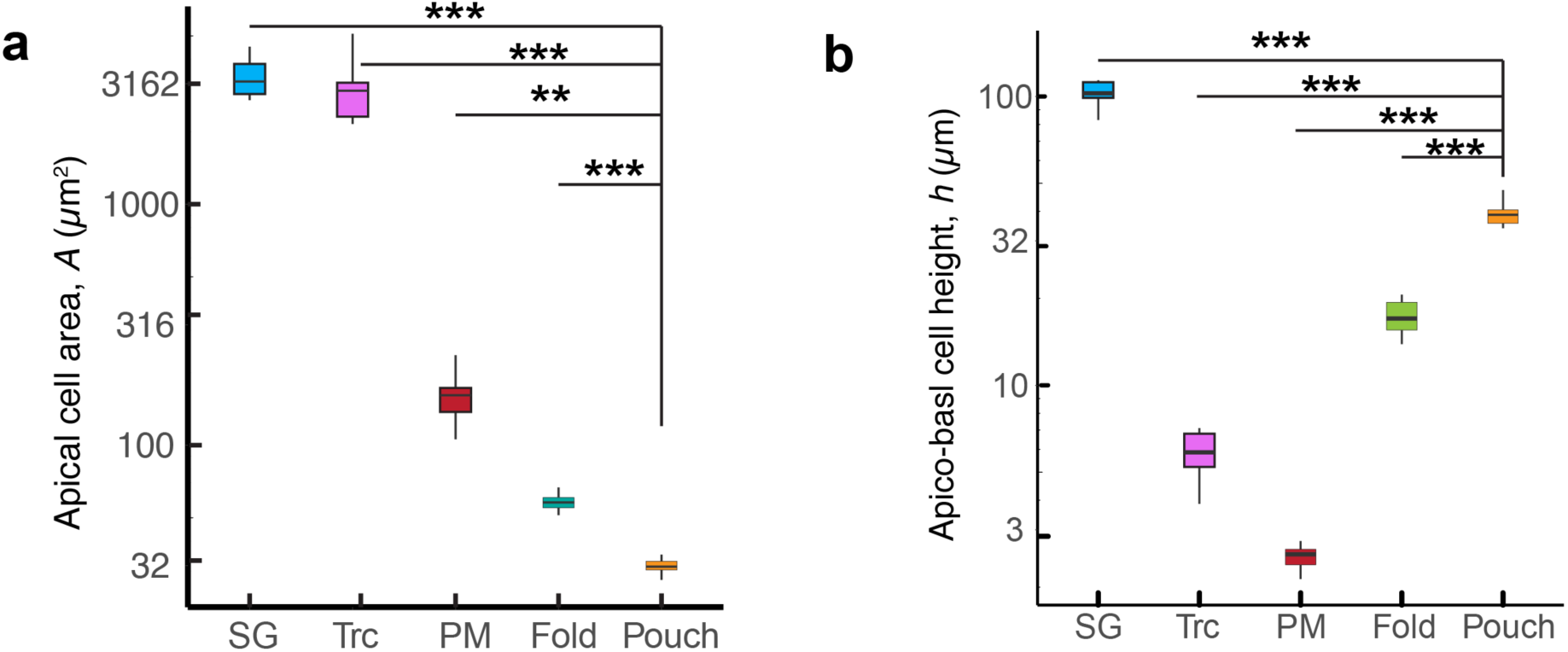
Apical cell area and apico-basal height in Drosophila tissues. **(a)** Box plot showing apical cell area in *Drosophila* tissues. Y -axis is shown in logscale. **(b)** Box plot showing apico-basal cell height in *Drosophila* tissues. Y-axis is shown in logscale.

**Supplementary Fig. 3:**
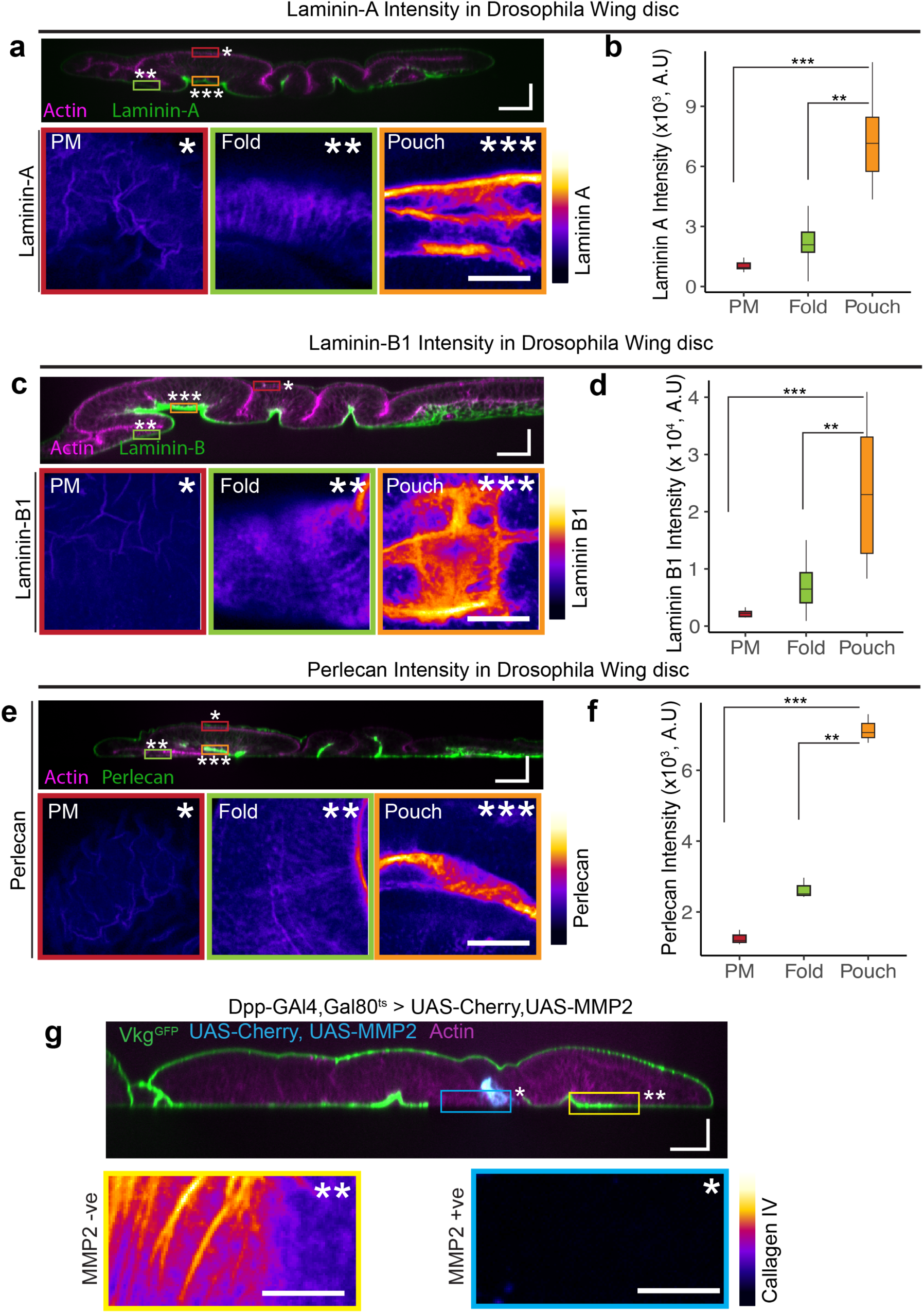
ECM protein concentrations in *Drosophila* wing disc. **(a)** Top panel: Cross section view of the wing disc expressing Laminin-A::GFP (green). Actin is shown in magenta. Bottom panel: Enlarged XY sections of the regions marked by yellow ROI (PM), green ROI (fold) and red ROI (pouch). Images show color coded Laminin-A intensity. **(b)** Box plot showing Laminin-A intensity in different regions of the wing disc. **(c)** Top panel: Cross section view of the wing disc expressing Laminin-B1::GFP. Actin is shown in magenta. Bottom panel: Enlarged XY sections of the regions marked by yellow ROI (PM), green ROI (fold) and red ROI (pouch). Images show color coded Laminin-B1 intensity. **(d)** Box plot showing Laminin-B1 intensity in different regions of the wing disc. **(e)** Top panel: Cross section view of the wing disc expressing Perlecan::GFP. Actin is shown in magenta. Bottom panel: Enlarged XY sections of the regions marked by yellow ROI (PM), green ROI (fold) and red ROI (pouch). Images show color coded Perlecan intensity. **(f)** Box plot showing Perlecan intensity in different regions of the wing disc. **(g)** Top panel: Cross-section view of the wing disc showing vkgGFP (Collagen IV) in green, actin in magenta and UAS-MMP2,UAS-CD8-Cherry in blue. MMP2 in the wing disc is driven by Dpp-GAL4,Gal80^ts^. Bottom panel: Enlarged XY sections Collagen IV of the regions marked by blue ROI (MMP2 +ve) and yellow ROI (MMP2 -ve). Images are color coded for Collagen IV intensity. Scale bar in the cross-section images, 25 µm along both axes. Scale bar in enlarged images in (a,c,e), 25 µm. Scale bar in enlarged images in (g), 15 µm. P-values are estimated using Student’s t-test. ** represents p < 0.001 and *** represents p < 10^−5^.

**Supplementary Fig. 4:**
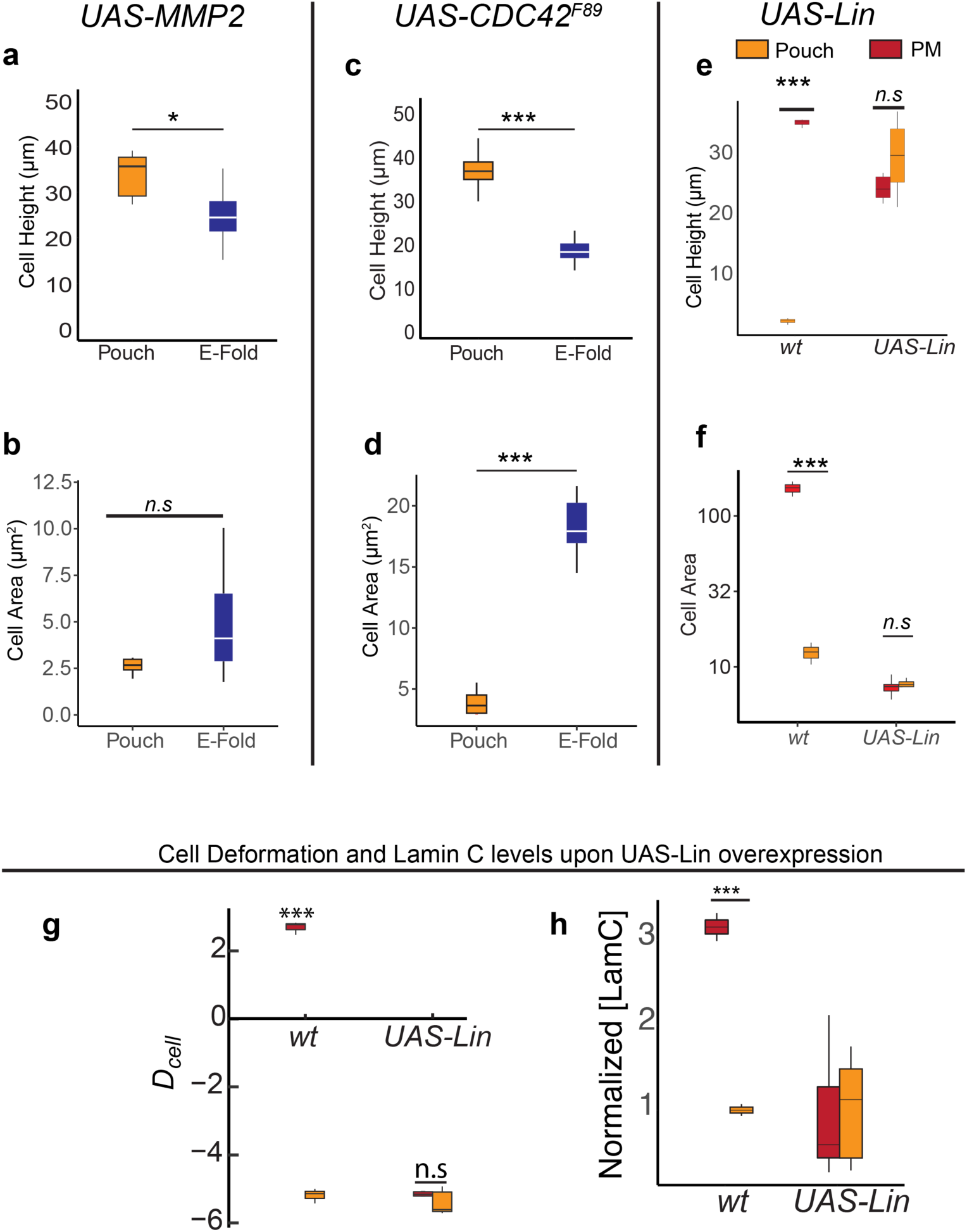
Cell height and cell area in *Drosophila* wing disc under different perturbations. **(a-b)** Cell height (a) and cell area (b) in the pouch (orange) and ectopic fold (green) in wing disc expressing UAS-MMP2 by *Dpp-Gal4,Gal80*^*ts*^. **(c-d)** Cell height (c) and cell area (d) in the pouch (orange) and ectopic fold (green) in wing disc expressing UAS-CDC42^F89^ by *Dpp-Gal4,Gal80*^*ts*^. **(e-f)** Cell height (e) and cell area (f) in PM (red) and pouch (blue) regions of the wing disc expressing either w- or *Lin* by Ubx-Gal4. **(g)** Box plot showing *D*_*cell*_ in wt and *UAS-Lin* overexpression in pouch and PM. **(h)** Box plot showing normalized LamC in wt and *UAS-Lin* overexpression in pouch (blue) and PM (red). Normalization is done with respect to the pouch. P-values are estimated using Student’s t-test. * represents p < 0.05, *** represents p < 10^−5^ and n.s represents that the differences are not statistically significant.

**Supplementary Fig. 5:**
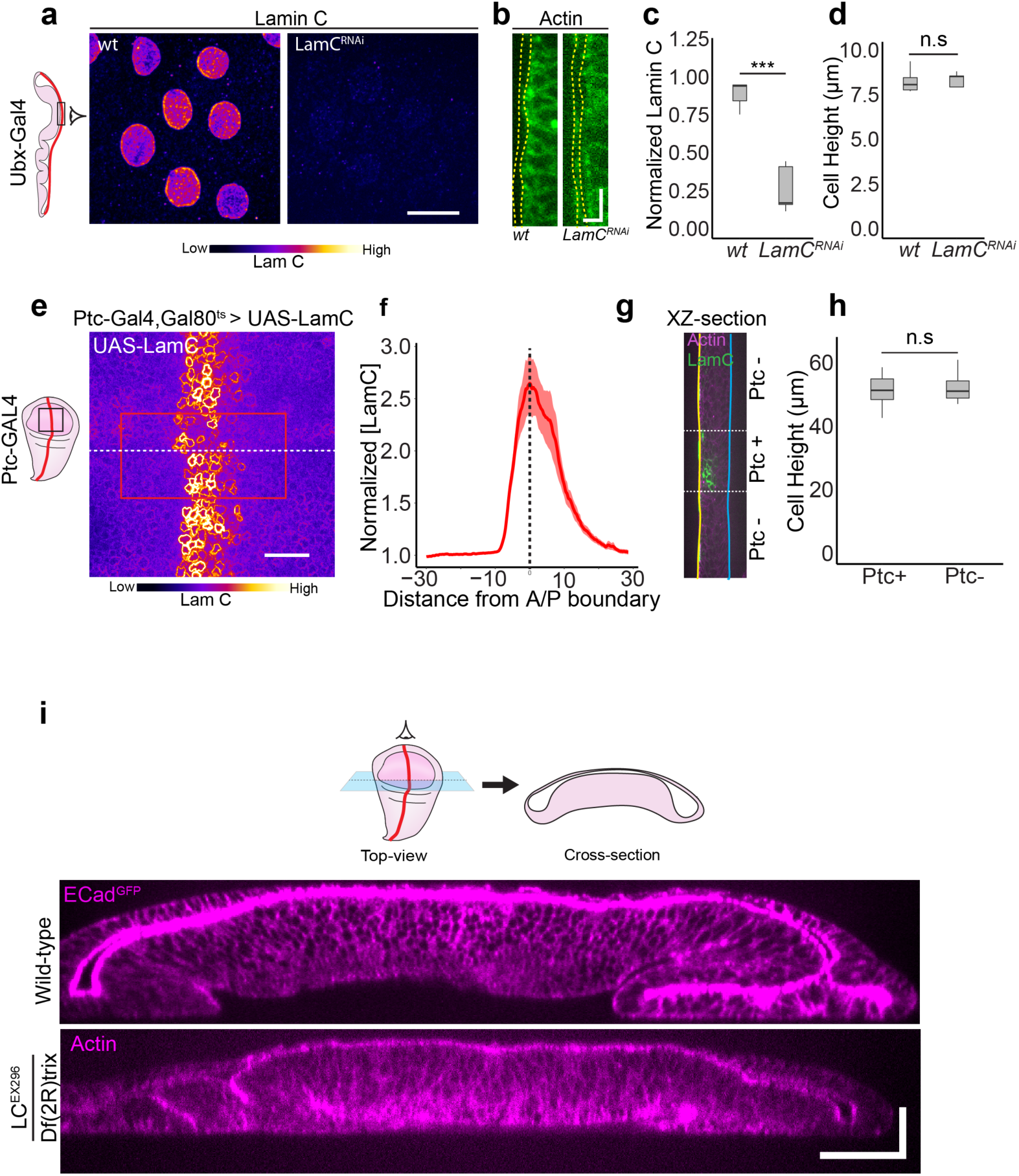
Morphology of wing disc cells is independent of Lamin C expression. **(a)** Images showing LamC in the PM region of the wing discs, in which Ubx-Gal4 either drives w- (left) or UAS-LamC^RNAi^(right). Scale bar, 15 µm **(b)** Cross-section views of the PM region of the wing disc shown actin (in green) for discs driving w- (left) or UAS-LamC^RNAi^ (right). Yellow dotted lines show the width of the PM layer of the wing disc. Scale bar, 10 µm (vertical), 5 µm (horizontal) **(c)** Box plot showing normalized LamC in the PM region of the wing disc. Normalization is performed w.r.t wt. **(d)** Box plot showing cell height in w- and LamC^RNAi^ expressing cells. **(e)** Image shows LamC in the pouch region of the wing disc, where LamC is overexpressed by Ptc-GAL4 stripe in temperature controlled manner using Gal80^ts^. The image is color coded for LamC intensity. Scale bar, 15 µm.**(f)** Normalized LamC intensity profile for the region marked by red ROI in (e). The mean over the region is plotted by the solid red line and the shaded region in red shows the standard deviation. Vertical dotted line marks the A/P boundary. **(g)** XZ section of the wing disc along the dotted line shown in (e). Actin is shown in magenta and LamC is shown in green. The yellow and blue solid lines mark the apical and basal surface of the wing pouch respectively. The two dotted lines marks the region of Ptc expression, where LamC is overexpressed. **(h)** Box plot showing cell height in Ptc+ and Ptc-cells. (**i**) Cross-section view of wild-type wing disc, and wing disc mutant for LamC (LamC^EX296^/Df(2R)trix. Top panel: wild-type disc with ECadGFP labelled in magenta. Bottom panel: Mutant disc with Actin labelled in magenta. Scale bar along both axes, 25 µm. P-values are estimated using Student’s t-test. *** represents p<10^−5^ and n.s represents that differences are not statistically significant.

**Supplementary fig. 6:**
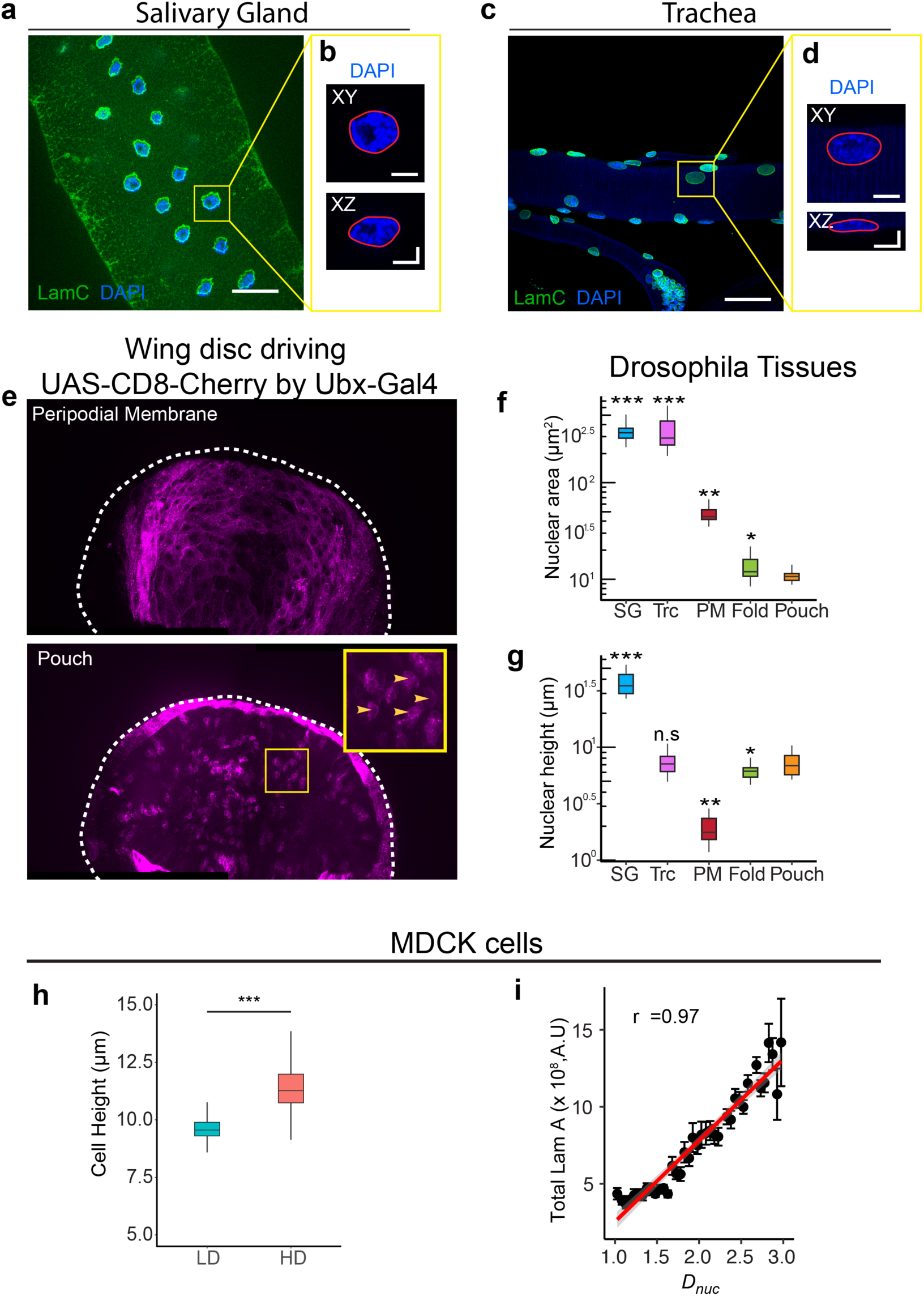
Nuclear morphology in Epithelial tissues. **(a)** Image showing LamC (green) and DAPI (blue) in Salivary gland. Scale bar, 50 µm **(b)** Enlarged image of the region marked by Yellow ROI in (a). Top panel: XY image and bottom panel: XZ cross-section image. Nucleus is labelled by DAPI in blue and the nuclear outline is marked by solid red line. Scale bar, 10 µm along both axes. **(c)** Image showing LamC (green) and DAPI (blue) in Trachea. Scale bar, 50 µm **(d)** Enlarged image of the region marked by yellow ROI in (a). Top panel: XY image and bottom panel: XZ cross-section image. Nucleus is labelled by DAPI in blue and the nuclear outline is marked by solid red line. Scale bar, 10 µm along both axes. **(e)** Wing disc driving UAS-CD8-Cherry by Ubx-Gal4. Left panel shows the PM region and right panel shows the pouch region. Inset image in the right panel is the enlarged image of the region marked by yellow ROI. Arrowheads show individual cells labelled by UAS-CD8-Cherry. Scale bar, 50 µm. **(f)** Box plot showing ox nuclear cross-sectional area in different *Drosophila* tissues. Y axis is shown in logscale. **(g)** Box plot showing nuclear height in different *Drosophila* tissues. Y axis is shown in logscale. (**h**) Box plot showing cell height in low density and high density cultures.**(i**) Binned scatter plot between *D*_*nuc*_ and total Lamin A in MDCK cells. Solid red line shows the linear fit to the data. P-values were estimated using one-way ANOVA and the comparison is shown w.r.t the wing pouch. * represents p < 0.05, ** represents p < 0.0001, *** represents p < 10^−5^, n.s represents that the differences are not statistically significant.

